# ESCRT-III/Vps4 controls heterochromatin-nuclear envelope attachments

**DOI:** 10.1101/579805

**Authors:** Gerard H. Pieper, Simon Sprenger, David Teis, Snezhana Oliferenko

## Abstract

In eukaryotes chromosomes are compartmentalized within the nucleus delimited by the double membrane of the nuclear envelope (NE). Defects in the function and structure of the NE are linked to disease^1,2^. During interphase, the NE organizes the genome and regulates its expression^3^. As cells enter mitosis, chromosomes are released from the NE, which is then remodelled to form the daughter nuclei at mitotic exit^4^. Interactions between the NE and chromatin underpinning both interphase and post-mitotic NE functions are executed by inner nuclear membrane (INM) proteins such as members of the evolutionarily conserved chromatin-binding LEM-domain family^5–8^. How chromatin tethering by these transmembrane proteins is controlled in interphase and if such a regulation contributes to subsequent NE dynamics in mitosis remains unclear. Here we probe these fundamental questions using an emerging model organism, the fission yeast *Schizosaccharomyces japonicus*, which breaks and reforms the NE during mitosis^9,10^. We show that attachments between heterochromatin and the transmembrane Lem2-Nur1 complex are continuously remodelled in interphase by the ESCRT-III/AAA-ATPase Vps4 machinery. ESCRT-III/Vps4 mediates the release of Lem2-Nur1 from heterochromatin as a prerequisite for the timely progression through mitosis. Failure in this process leads to persistent association of chromosomes with the INM, which prevents Lem2-Nur1 from re-localizing to the sites of NE sealing around the mitotic spindle and severely delays re-establishment of nucleocytoplasmic compartmentalization. Our work establishes the INM transmembrane Lem2-Nur1 complex as a ‘substrate’ for ESCRT-III/Vps4 to couple dynamic tethering of chromosomes to the INM with the establishment of nuclear compartmentalization.

---

In *S. japonicus*, the NE ruptures in anaphase at a single medial site, to be quickly reformed around the segregated genomes at mitotic exit^10,11^. The establishment of nucleocytoplasmic compartmentalization begins while the spindle is still present^10^. At this stage, the nuclear membrane wraps tightly around the intersecting spindle microtubules^10,11^. Thus, nuclear compartmentalization is established prior to nuclear membrane resealing, which can be completed only after spindle breakdown. Lem2 enriches at these structures that we term ‘tails’, in addition to its spindle pole body (SPB) localization^10^ (Fig. 1a). Other NE proteins such as components of the nuclear pore complexes (NPCs) and the second fission yeast LEM-domain protein Man1 are largely excluded from the ‘tails’ due to their interactions with segregating chromosomes at this stage of mitosis^10–12^. Thus, the ‘tail’ is a specialized membrane domain, spatially segregated and partitioned away from the rest of the NE during mitotic exit (Fig. 1a, right panel).

**Figure 1.**
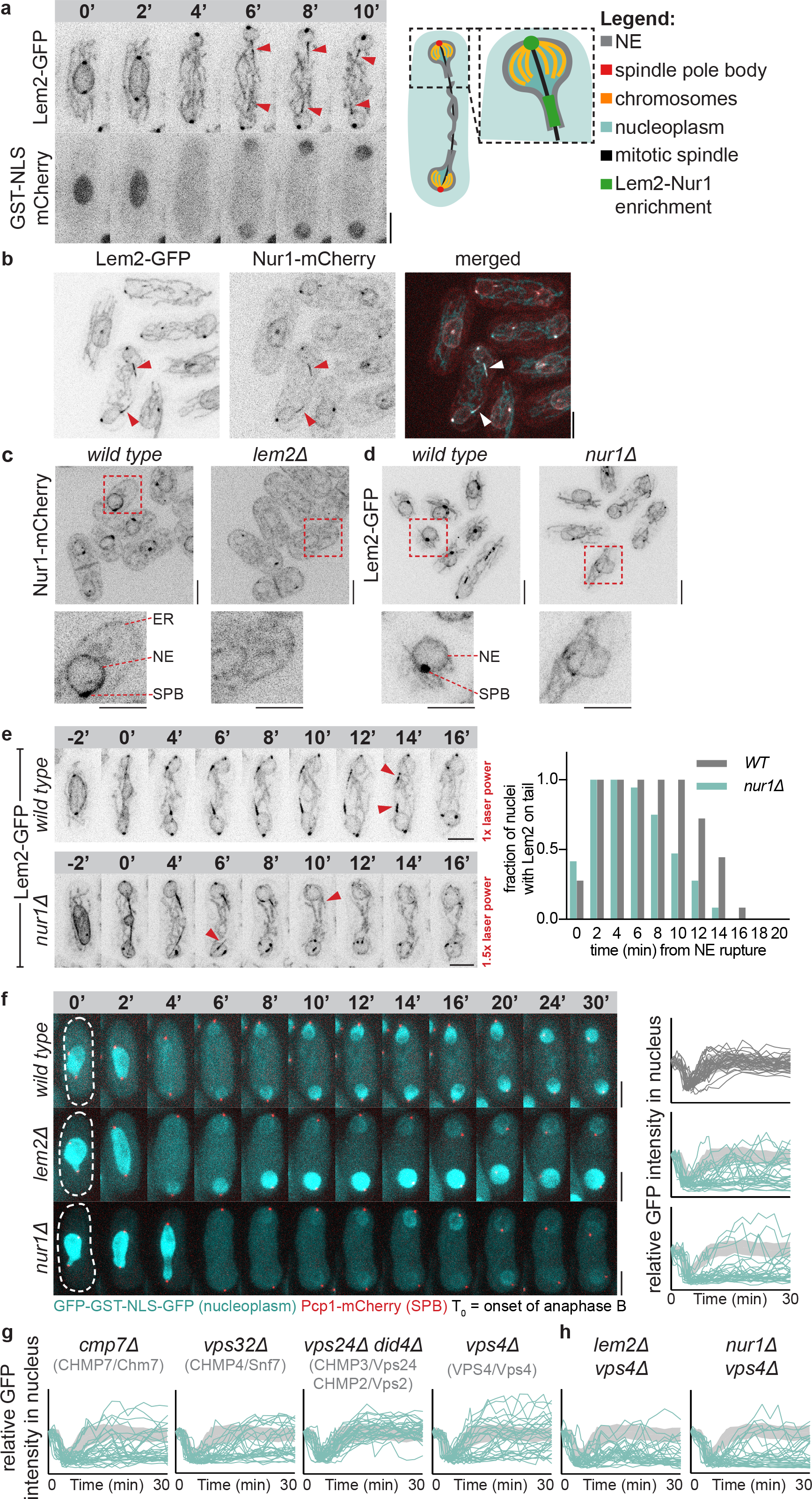
Lem2-Nur1 together with a minimal ESCRT-III/Vps4 machinery mediate NE resealing in *S. japonicus*. (**a**) Time-lapse of a representative WT *S. japonicus* cell expressing Lem2-GFP and GST-NLS-mCherry undergoing mitosis, starting from early anaphase B. Arrows indicate beginning of Lem2 enrichment at the ‘tails’, which coincides with establishment of nucleocytoplasmic compartmentalisation. Scheme represents the 8-minute time point. (**b**) Cells co-expressing Lem2-GFP and Nur1-mCherry. Arrows indicate their enrichment at the ‘tails’. (**c**) Nur1-mCherry expressing cells of indicated genotypes imaged and presented at the same settings for comparison of signal strength. See magnified images below. Shown are Z-projections of the middle 8 (out of 10) Z-slices over a distance of 3.5 µm. (**d**) Z-projections of images of Lem2-GFP expressing cells of indicated genotypes imaged and presented at the same settings for comparison of signal strength. (**e**) *Left panel*, Cells of indicated genotypes expressing Lem2-GFP. The *nur1∆* mutants were imaged using higher laser power (1.5x) as compared to the WT. Arrows indicate the last time point before Lem2 disappearance from the ‘tail’. *Right panel*, a graph representing the duration of Lem2 presence on the ‘tails’ in cells of indicated genotypes. (**f**) NE resealing assay in cells of indicated genotypes co-expressing GFP-NLS and Pcp1-mCherry. Time-lapse sequences were obtained throughout mitosis. Quantification of the GFP fluorescence intensity in the nucleus is relative to the time point 0, prior to NE breakage (n= 15 cells / 30 nuclei). The standard deviation of the WT quantification for each time-point (light grey) is shown together with the quantification of each mutant to aid comparison. (**g**) NE resealing assay for ESCRT-III/Vps4 mutants performed as in (**f**). See representative cells in Supplemental Fig. 1a. (**h**) NE resealing assay performed for *lem2∆ vps4∆* and *nur1∆ vps4∆* double mutants as in (**f**). See representative cells in Supplemental Fig. 1b. (**a**, **b**, **d**, **e**, **f**). Shown are Z-projections of spinning disk confocal stacks. Scale bars represent 5 μm.

Lem2 orthologues function in complex with another INM protein Nur1 to promote heterochromatin tethering and heterochromatic gene silencing^13–15^. In *S. japonicus*, Nur1 largely co-localized with Lem2 throughout the cell cycle, enriching at the NE ‘tails’ at the end of mitosis (Fig. 1b). This localization of Nur1 required Lem2, as Nur1 redistributed throughout the ER in *lem2∆* cells (Fig. 1c). Conversely, in cells lacking Nur1, less Lem2 signal was detected at the SPBs and the NE (Fig. 1d). Additionally, Lem2 residence time at the ‘tails’ was drastically reduced in Nur1-defcient cells (Fig. 1e). Thus, Lem2 and Nur1 appear to co-depend on each other for proper localization to the NE and the SPB throughout the cell cycle.

We have previously shown that proper post-mitotic establishment of nucleocytoplasmic compartmentalisation failed in the absence of Lem2^10^ (Fig. 1f). Using a nucleoplasmic reporter protein GFP-GST-NLS-GFP (referred to as GFP-NLS), we discovered that *nur1∆* cells displayed a similar phenotype. Compared to wild type (WT) cells where nucleocytoplasmic compartmentalisation was typically re-established 4-6 minutes after NE rupture in anaphase, *lem2∆* and *nur1∆* mutants achieved this state considerably later and in a desynchronised manner. These mutants also frequently failed to maintain nuclear integrity after a seemingly successful recovery event (Fig. 1f). Following an extended delay, most mutant cells eventually recovered nuclear integrity. We concluded that the persistent enrichment of the Lem2-Nur1 complex at the sites where the NE wraps around the mitotic spindle might be essential for the timely re-establishment of nucleocytoplasmic compartmentalisation.

LEM-domain proteins were proposed to mediate NPC quality control in *S. cerevisiae*^16,17^, maintain NE integrity in *S. pombe*^18^, and help reform the NE in cultured human cells^18–20^ through recruitment of the ESCRT-III machinery^19,20^. Therefore, we systematically addressed if ESCRT-III/Vps4 contributed to establishing nucleocytoplasmic compartmentalisation in *S. japonicus*. Cells lacking the nuclear ESCRT-III adaptor Cmp7^20^ (*S. c*.: Chm7, *H. s*.: CHMP7), the major ESCRT-III subunit Vps32^21^ (*S. c*.: Snf7, *H. s*.: CHMP4) and the AAA-ATPase Vps4^22^ exhibited defects in re-establishing nucleocytoplasmic compartmentalisation following mitosis, similar to *lem2∆* or *nur1∆* cells (Fig. 1g, Extended Data Fig. 1a). The Lem2-Nur1 complex and ESCRT-III/Vps4 appeared to function in the same pathway since *lem2∆ vps4∆* and *nur1∆ vps4∆* double mutant cells exhibited phenotypes comparable to that of the single mutants (Fig. 1h, Extended Data Fig. 1b). Surprisingly, the core ESCRT-III subunits, Vps24 (*S. c.:* Vps24, *H. s.:* CHMP3) and Did4 (*S. c.:* Vps2, *H. s.:* CHMP2) that are essential for endosomal ESCRT functions^23^, did not appear to be involved in nucleocytoplasmic compartmentalisation in *S. japonicus* (Fig. 1g, Extended Data Fig. 1c). Similarly, the endosome-specific ESCRT-III adaptor, Vps25 (ESCRT-II), was not required for this process (Extended Data Fig. 1d). Consistent with our earlier report of a premature loss of nucleocytoplasmic integrity in Lem2-deficient mitotic cells^10^, we observed leaking of the nucleoplasmic marker in a subset of ESCRT-III/Vps4 mutants already at the onset of anaphase spindle elongation (Extended Data Fig. 1e). The persistence of the mitotic spindle was not affected by the lack of Lem2, Vps24 or Vps4 (Extended Data Fig. 1f), suggesting that ESCRT-III/Vps4-mediated establishment of nuclear compartmentalisation and spindle breakdown are not coupled. We concluded that in *S. japonicus*, the Lem2-Nur1 complex works together with ESCRT-III subunits Cmp7-Vps32 and the AAA-ATPase Vps4 to re-establish nuclear compartmentalisation prior to the disassembly of the spindle.

The vast majority of functional, endogenously tagged Vps4-3xHA-GFP and Vps24-LAP-GFP (Extended Data Fig. 2a; referred to as Vps4-GFP and Vps24-GFP) was detected on cytoplasmic objects adjacent to FM4-64-marked vacuoles (Fig. 2a, Extended Data Fig. 2b), reminiscent of multivesicular bodies (MVBs) in budding yeast^24^. Consistent with the endosomal nature of these perivacuolar objects, deletion of *vps27* (ESCRT-0), *vps28* (ESCRT-I), *vps25* (ESCRT-II), *vps24* or *did4* (ESCRT-III) resulted in re-localisation of Vps4-GFP into the cytoplasm (Extended Data Fig. 2b). Only a few Vps4-GFP objects remained and the majority appeared to be distinct from perivacuolar MVBs. Since the vast majority of Vps4-GFP localized to endosomes, it was difficult to detect Vps4 at the NE. Nevertheless, we observed Vps4-GFP localizing transiently to the Lem2 ‘tails’ and the SPBs during mitotic exit (Fig. 2a), the latter likely coinciding with SPB extrusion from the NE plane^25,26^. We confirmed this localization pattern by using a *vps25∆* mutant where Vps4 could no longer localize to endosomes (Fig. 2b and Extended Data Fig. 2b). Using this system, we established that Vps4-GFP was primarily recruited to the distal ends of the Lem2-mCherry-labelled NE ‘tails’ (Fig. 2b,c). Peak Vps4-GFP recruitment occurred around 7 min after NE rupture (Fig. 2d), concurrent with re-establishment of nuclear compartmentalisation (Fig. 1a).

**Figure 2.**
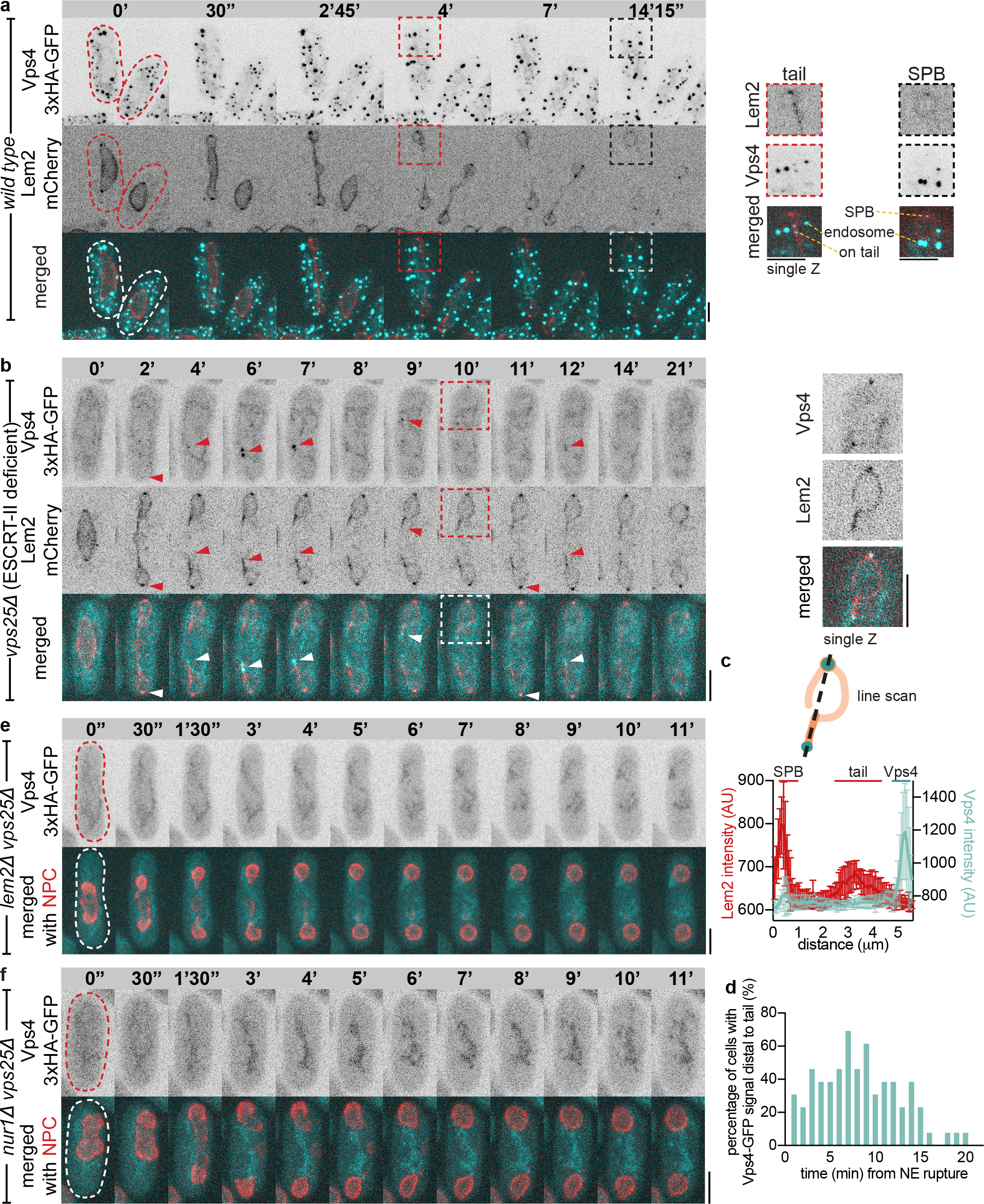
Lem2 together with Nur1 recruit ESCRT-III/Vps4 to the sites of NE sealing around the mitotic spindle. (**a**) Time-lapse maximum projection sequences of Vps4-GFP Lem2-mCherry expressing cells starting from prior to NE breakage. Magnified images focus on a single confocal slice showing Vps4 localisation to the SPB and to the ‘tail’. Note that the endosomal Vps4 signal drowns out the NE-localized Vps4. (**b**) A time-lapse Z-projected sequence of a *vps25∆* Vps4-GFP-Lem2-mCherry-expressing cell starting prior to NE rupture (n=21 cells). Arrows indicate Vps4 localisation to the SPBs and to the distal end of the ‘tail’ structure. A single Z-slice of an area outlined at a 10-minute time point is shown as a magnified image on the right. (**c**) Quantification of Vps4 enrichment at the distal end of the Lem2 tail (n=10). (**d**) Quantification of the timing of Vps4 recruitment to the ‘tails’, shown as a percentage of cells exhibiting Vps4-GFP signal. Cells were followed between NE rupture and spindle breakdown (n=13). (**e**) A time-lapse sequence of a *lem2∆ vps25∆* Vps4-GFP Nup189-mCherry cell starting prior to NE rupture (n=11). (**f**) A time-lapse sequence of a *nur1∆ vps25∆* Vps4-GFP Nup189-mCherry cell starting prior to NE rupture (n=17). (**a**, **b**, **e**, **f**) Shown are Z-projections of spinning disk confocal stacks. Scale bars represent 5 μm.

Vps4 recruitment to the distal ends of Lem2 ‘tails’ during mitotic exit depended on the functional Lem2-Nur1 complex. In the absence of Lem2 or Nur1 Vps4 was no longer detected at the NE (Fig. 2e,f and Extended Data Fig. 3a). Vps4 recruitment also depended on Cmp7 and Vps32 (Extended Data Fig. 3b,c). Vps24-GFP exhibited similar localization at mitotic exit, despite not being essential for re-establishment of nuclear integrity following NE breakdown (Extended Data Fig. 3d-e). Such a behaviour is evocative of dynamic ESCRT-III/Vps4 assemblies on endosomes^24,27^ and during HIV budding^28,29^. The timing of Vps4 recruitment that occurs throughout the lifetime of the spindle suggests that ESCRT-III/Vps4 is required to establish nucleocytoplasmic compartmentalisation prior to spindle disassembly and membrane resealing. At this stage ESCRT-III/Vps4 may play a role in the maintenance of the Lem2-Nur1 ‘tails’. Of note, in human cells ESCRT-III is also recruited to the nuclear membrane in close vicinity of spindle remnants^20^, before they are dismantled allowing membrane closure.

We set up an *in vitro* binding assay using *S. japonicus* proteins purified from *Escherichia coli* to analyse the interactions between Lem2 and the ‘minimal’ ESCRT-III/Vps4 components that were essential for nuclear re-compartmentalization. Therefore we purified the C-terminal nucleoplasmic domain (aa564-673) of Lem2 (GST-Lem2^564-673^) encompassing the MAN1-Src1-C-terminal (MSC) domain^18^, the C-terminal ESCRT-III-like domain (aa 242-436) of Cmp7^242-436^-3xFLAG, Vps32-3xMYC as well as Vps4-3xHA and an ATP hydrolysis deficient Vps4^E233Q^3xHA substrate trap mutant that binds ATP but cannot hydrolyse it (Vps4^EQ^)^22^. Purified Did4 was used as a positive control for Vps4 binding^30,31^ (Extended Data Fig. 4a).

The substrate trap Vps4^EQ^ interacted directly with Cmp7^242-436^ and Vps32 but not with Lem2^564-673^, suggesting that either ESCRT-III protein could recruit Vps4 to Lem2 (Extended Data Fig. 4a). Lem2^564-673^ interacted directly with Cmp7^242-436^. Thus, the C-terminal MSC domain of Lem2 is sufficient for binding to the C-terminal ESCRT-III-like domain of Cmp7 (Fig. 3a, lane 8). Direct binding of Vps32 to Lem2^564-673^ was not detected (Fig. 3a; lane 10). Yet, Vps32 was capable of associating with preformed Lem2^564-673^ - Cmp7^242-436^ complexes (Fig. 3a; lane 11). When Vps4 was added to these preformed complexes, it released Vps32 and Cmp7^242-436^ from Lem2^564-673^ (Fig. 3a; lanes 9 and 12), in an ATP-dependent manner (Extended Data Fig. 4b; lane 5). In contrast, the substrate trap mutant Vps4^EQ^ failed to disassemble Lem2^564-673^ - Cmp7^242-436^ interaction and instead, was retained on these complexes (Fig. 3b; lane 9). Significantly more Vps4^EQ^ was trapped in complexes containing Vps32 in addition to Cmp7^242-436^ (Fig. 3b; lane 12). Collectively, these results suggested that Vps4 could disassemble the interaction between Cmp7 and Lem2 *in vitro* (Fig. 3c).

**Fig. 3.**
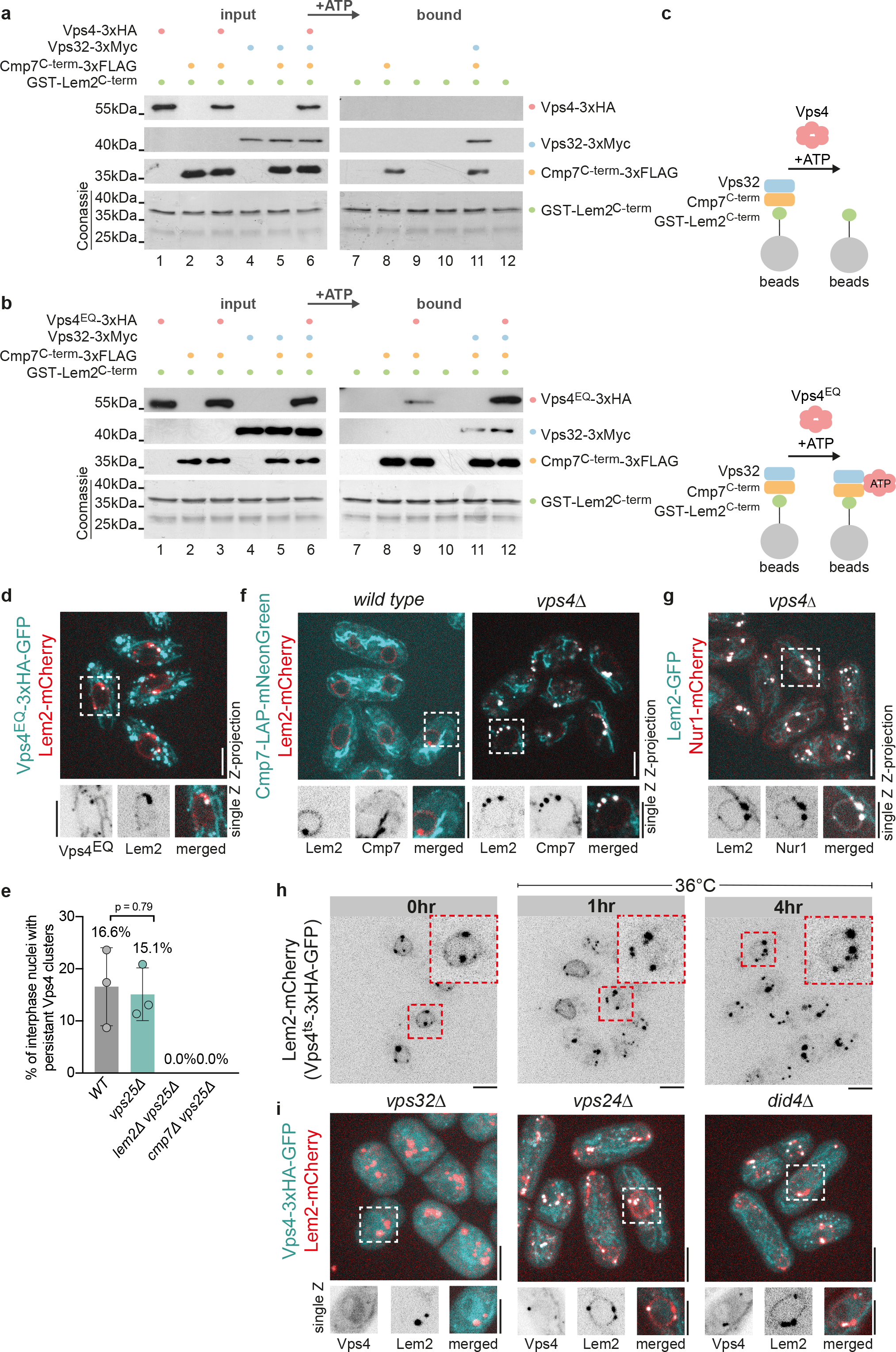
Vps4 disassembles Lem2-Cmp7 interactions. (**a**) *In vitro* binding assay. GST-Lem2^C-term^ captured at glutathione beads was first incubated with Cmp7^C-term^ with or without Vps32 followed by wash-out of the unbound protein. This complex was then incubated with Vps4. Binding experiments were performed in the presence of ATP. The left panel shows the protein input with Coomassie staining for Lem2^C-term^ and by Western-blotting for ESCRT-III/Vps4. The right panel shows the proteins recovered from Lem2-C-term-bound beads after the binding reactions. (**b**) *In vitro* binding assay with the same layout as in (**a**), but with the substrate trap Vps4EQ mutant added instead of the wild type Vps4. Note that Vps4EQ becomes locked onto beads containing Lem2-Cmp7 or Lem2-Cmp7-Vps32 (lane 9 and 12). (**c**) A schematic overview of experimental setup and results of *in vitro* binding assays shown in (**a**) and (**b**). (**d**) A maximum projection of a spinning disk confocal stack of interphase cells expressing Vps4^EQ^-GFP and Lem2-mCherry. Below see a magnified single slice image focussing on colocalisation of Vps4^EQ^ with Lem2 in clusters at the NE. (**e**) A graph showing the percentage of interphase cells of indicated genotypes harbouring a persistent Vps4 focus at the NE (n=3 experiments with at least 40 cells counted for each experiment; error bars indicate standard deviation; T-test). Note that no persistent clusters were observed in *lem2∆* or *cmp7∆* mutants. (**f**) Maximum projections of spinning disk confocal stacks of WT and *vps4∆* cells expressing Cmp7-LAP-mNeonGreen and Lem2-mCherry. See a magnified single slice image showing a ER localisation of Cmp7 and NE/SPB localisation of Lem2. Note that Cmp7 could only be faintly detected on the cortical ER; these images show high mitochondrial autofluorescence because of very weak specific signal. In *vps4∆* cells Cmp7 becomes clustered at the NE together with Lem2. (**g**) A maximum projection of a spinning disk confocal stack of *vps4∆* cells expressing Lem2-GFP and Nur1-mCherry. (**h**) A time course of cells expressing Lem2-mCherry in the temperature sensitive Vps4^ts^ background. Cells grown at 25°C were shifted to 36°C to inactivate Vps4^ts^ and samples were taken at 1 and 4 hours (n=2 experiments, at least 30 cells for each experiment per time-point). (**i**) Maximum projection images of spinning disk confocal stacks of cells of indicated genotypes co-expressing Vps4-GFP and Lem2-mCherry. (**d**, **f**, **g, h, i**) Scale bars represent 5 μm.

Consistent with this model, expression of Vps4^EQ^-GFP in live cells trapped it together with Lem2 at the NE (Fig. 3d). In addition, Vps4^EQ^-GFP was detected on endosomes. Confirming that these interactions also occur in WT cells, we observed persistent NE-associated foci of Vps4-GFP in a fraction of WT interphase cells (Fig. 3e; Extended Data Fig. 4c; typically, one Vps4-GFP object per nucleus). The frequency of these foci was not affected by the deletion of *vps25* but they were no longer present when *lem2* or *cmp7* were deleted (Fig. 3e). In the absence of Vps4, Cmp7-mNeonGreen was trapped in large clusters at the NE together with Lem2-Nur1 (Fig. 3f,g). In WT cells Cmp7-mNeonGreen was faintly visible around the cell cortex and sometimes at the NE (Fig. 3f), indicating its ER localisation as previously observed for the budding yeast and human Cmp7 orthologues^32,33^. To analyse how Lem2 clusters arise, we acutely inactivated Vps4, using a temperature sensitive mutant, Vps4^ts^-GFP (*vps4-*^I307T,L327S^)^22^. At 25°C, Lem2 remained largely distributed around the NE and the SPBs, although we detected a few Lem2 clusters at the NE, suggesting that Vps4^ts^-GFP behaved as a mild hypomorph (Fig. 3h). Upon the shift to the restrictive temperature of 36°C, Lem2 became increasingly trapped in clusters at the NE (Fig. 3h). This suggested that continuous Vps4 activity is required to prevent the formation of persistent Lem2 clusters at the INM during interphase.

All core ESCRT-III mutants exhibited Lem2 clustering. Yet, the severity of this phenotype depended on whether Vps4 was present in these clusters and correlated with failed nuclear compartmentalization (Fig. 1g). Similar to *vps4∆* cells, Lem2 was found exclusively in large clusters in *vps32∆* mutants (Fig. 3i). Vps4 did not localize to these clusters, suggesting that *in vivo*, unlike *in vitro*, Vps32 was essential to recruit Vps4 to Lem2 and Cmp7. In the absence of Vps24 and Did4, Lem2 clusters were smaller in size and contained Vps4 (Fig. 3i). Thus, in these mutants Vps4 could be at least partially functional and disassemble Lem2-Cmp7-Vps32 complexes at the NE. As expected, Lem2 localization was not affected by the loss of ESCRT-0, -I and –II function (Extended Data Fig. 4d). Taken together, our data suggest that the ESCRT-III/Vps4 machinery prevents aberrant clustering of Lem2-Nur1 complexes at the NE.

Since the Lem2-Nur1 complex is involved in heterochromatin maintenance and its tethering to the NE^13,14,34,35^, we analysed if the Nur1-Lem2 clusters in ESCRT-III-deficient cells co-localized with heterochromatin. In fission yeast, all heterochromatin including sub-telomeric and peri-centromeric sequences localises to the nuclear periphery during interphase^36^. Consistently, Lem2-GFP clusters were adjacent to heterochromatic domains at the INM marked by the heterochromatin protein 1 (HP1) orthologue Swi6-mCherry^37^ (Fig. 4a). *S. japonicus*, like other eukaryotes, releases chromosomes from the INM as it enters mitosis^4,38–41^. Consistent with this, Lem2 clusters resolved during mitosis in *cmp7∆* and *vps24∆* cells. Remarkably, this was not the case in *vps32∆* and *vps4∆* mutants where Lem2-heterochromatin clusters persisted throughout mitosis (Fig. 4a). This indicated that Lem2-heterochromatin attachments in these genetic backgrounds were refractory to mitotic regulation responsible for chromatin release^4^. Such severe association of Lem2 with heterochromatin in *vps4∆* mutant cells was prevented in the absence of Nur1, indicating that Nur1 linked Lem2 to heterochromatin (Fig. 4b and Extended Data Fig. 5a). De-clustering of Lem2 during mitosis in *vps24∆* cells is likely due by the fact that Vps4 is still recruited to the NE in the absence of Vps24 and is capable of residual function (Fig. 3i). Yet, we have shown that Vps4 is not recruited to the NE in the absence of Cmp7 (Extended Data Fig. 3b and Fig. 3e). We interpret this as evidence that Cmp7 plays an integral role in driving the formation of persistent heterochromatin-associated Lem2-Nur1 clusters in the absence of Vps4 function.

**Fig. 4.**
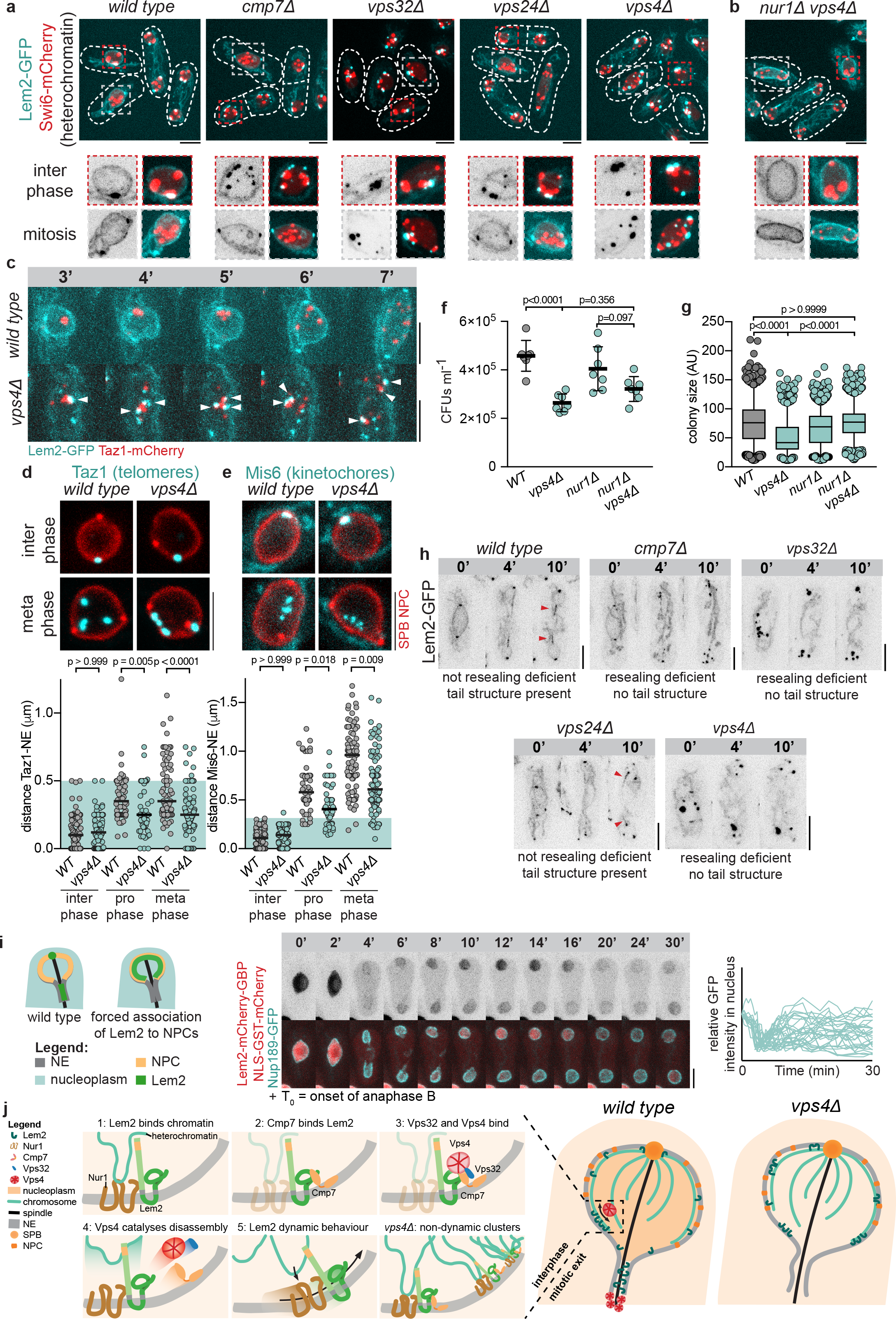
ESCRT-III/Vps4 is required for remodelling heterochromatin-Lem2 association in interphase and subsequent relocalization of Lem2 to the sites of NE sealing at mitotic exit. (**a**) *Top panel*, maximum projections of spinning disk confocal stacks of cells of indicated genotypes co-expressing Lem2-GFP and Swi6-mCherry. *Bottom panel*, an interphase and a mitotic cell are highlighted in magnified images. Note that in *S. japonicus*, unlike in *S. pombe*, Swi6 associates with chromosomes throughout mitosis. (**b**) Same set-up as for (**a**) but for *nur1∆ vps4∆* mutant cells. (**c**) Time-lapse maximum projection sequences of WT (top) and *vps4∆* (bottom) cells expressing Lem2-GFP Taz1-mCherry, focusing on mitotic nuclei. For full time courses and images of entire cells see Supplemental Fig. 5b. Arrows indicate the fate of a single Lem2-associated telomere cluster that splits and the separated telomeres that stay associated with Lem2 throughout mitosis. (**d**) Measurements of Taz1-NE distance at indicated stages of the cell cycle. Z-projections of middle 8 confocal slices showing the nuclei of the WT and *vps4∆* Taz1-GFP Nup189-mCherry Pcp1-mCherry expressing cells (*top*). Quantification of Taz1-NE distance, as described in Materials and Methods. Shown in green is the full range of WT interphase data (*bottom*). Kruskal-Wallis test, Dunn’s test for multiple comparisons. (**e**) Same set-up as in (**d**) but for Mis6-NE distance. (**f**) Quantification of CFU assays for cells of indicated genotypes (n=7). One-way ANOVA, Tukey’s multiple comparisons test. (**g**) Quantification of colony size in the CFU assay shown in (**g**). Presented is the pooled data of 3 technical replicates of 3 biological experiments. Kruskal-Wallis test, Dunn’s test for multiple comparisons. (**h**) Maximum projections of time-lapse spinning disk confocal stacks of cells of indicated genotypes expressing Lem2-GFP starting from prior to NE breakdown. Arrows indicate the presence of ‘tail’ structures in the WT and *vps24∆* mutant. (**i**) Synthetic tethering of Lem2-mCherry-GBP to Nup189-GFP. *Left*, a scheme of the forced association of Lem2 to Nup189 that prevents its enrichment at the ‘tails’. *Middle*, a representative cell exhibiting a NE resealing phenotype. *Right*, quantifications of relative GFP intensity in the nuclei of 15 cells (30 nuclei), analysed as in Fig. 1. (**j**) A cartoon explaining our model for ESCRT-III/Vps4 function in interphase chromatin tethering to the NE and the resulting NE sealing phenotypes during mitosis. 1: Lem2 binds heterochromatin, for which Nur1 is required. 2: Cmp7 recognises the heterochromatin bound state of Lem2-Nur1. 3: Cmp7 then recruits Vps32 and the AAA-ATPase Vps4. 4: Vps4 now catalyses the disassembly of the Lem2-Cmp7 interaction. 5: This results in Lem2-Nur1 complex releasing heterochromatin and being free to move around the NE and re-engage heterochromatin in cycles of binding and unbinding. In the absence of Vps4 function the interaction between Lem2 and Cmp7 persists, resulting in the stable association of Lem2-Nur1 with Cmp7 and heterochromatin. During mitotic exit, such a stable engagement prevents Lem2 from enriching at the ‘tail’, which we hypothesise results in a destabilised interphase between membrane and the spindle and a delay in re-establishing the nucleocytoplasmic compartmentalisation. (**a**, **b**, **c**, **d**, **e**, **h**, **i**) Scale bars represent 5

We investigated the fate of persistent Lem2 clusters in mitotic *vps4∆* cells by performing time-lapse imaging of Lem2-GFP together with a telomere marker Taz1-mCherry that borders the sub-telomeric heterochromatin. In the WT, the NE-bound telomere clusters dispersed during early mitosis (Fig. 4c and Extended Data Fig. 5b) and moved away from the NE, indicating that chromosome arms were released^12,39^ (Fig. 4d). In *vps4∆* cells Taz1 remained associated with Lem2 clusters throughout mitosis, which co-segregated together with telomeres (Fig. 4c and Extended Data Fig. 5b). Indeed, most telomeres did not appear to move away from the NE in mitotic *vps4∆* cells as compared to the WT (Fig. 4d). Kinetochores of *vps4∆* cells labelled by Mis6-GFP did detach from the nuclear periphery, yet this distance was significantly reduced as compared to the WT (Fig. 4e), indicating that their movement was spatially constrained, likely as a consequence of failed telomere release. The persistent mitotic chromosome arm-NE association deregulated mitotic progression in *vps4∆* cells. In the WT, the spindle assembly checkpoint (SAC) protein Mad2-GFP^42^ typically remained on kinetochores for 5-10 minutes (Extended Data Fig. 5c). However, in *vps4∆* cells, we observed an abnormally broad distribution of the duration of SAC activation. Whereas some cells were delayed at the metaphase-to-anaphase transition, others had a marked decrease in duration of Mad2 kinetochore recruitment (Extended Data Fig. 5c).

The growth defect of *vps4∆* in *S. pombe* was rescued by the deletion of *lem2* or *cmp7*^18^, which in light of our results might be explained by the disruption of persistent chromosome-NE associations. Indeed, in the absence of Vps4 in *S. pombe*, Lem2 exhibited severe clustering throughout mitosis (Extended Data Fig. 5e), suggesting that ESCRT-III/Vps4 function in preventing Lem2-heterochromatin clustering is evolutionarily conserved. Similarly, the lack of Vps4 caused profound defects in viability of *S. japonicus* – *vps4∆* cultures exhibited significant decrease in the number and size of colonies when grown on solid medium (Fig. 4f,g and Extended Data Fig. 5d). The growth defects were at least partially associated with persistent chromatin-NE associations, since the colony size, although not the number of colony forming units was rescued when Lem2 clustering was prevented by additionally removing Nur1 (Fig. 4f,g; Extended Data Fig. 5d). This was despite the fact that the *nur1∆ vps4∆* double mutant was defective in nucleocytoplasmic compartmentalisation (Fig. 1h). We therefore concluded that persistent heterochromatin association with Lem2-Nur1 at the NE was a major cause of growth defects in Vps4-deficient cells.

We hypothesized that the function of ESCRT-III/Vps4 in releasing Lem2 from heterochromatin was required for Lem2 to enrich at the ‘tails’. Indeed, in the mutants that were deficient in nuclear re-compartmentalization (*nur1∆*, *cmp7∆*, *vps32∆* and *vps4∆*) Lem2 failed to enrich on ‘tail’ structures (Fig. 4h and Fig. 1e). In the *vps24∆* mutant that was not defective in this process, Lem2 was still present at the ‘tails’ (Fig. 4h). To formally test this hypothesis, we tagged Lem2 with GFP-binding-protein (GBP) and expressed it together with Nup189-GFP, forcing the association between Lem2 and the NPCs (Extended Data Fig. 5f). The NPCs are excluded from the ‘tail’ domain and instead are associated with chromosome arms through Man1^12^. Indeed, Lem2-GBP/Nup189-GFP complexes localized around the NE but were absent from ‘tails’ (Extended Data Fig. 5f). Failure to localize Lem2 to the regions where the spindle intersects with the NE led to a marked deficiency in re-establishing nucleocytoplasmic compartmentalisation (Fig. 4i). Under these conditions, ESCRT-III and Vps4 were fully functional, but Lem2 was sequestered away from the tails. Hence the function of ESCRT-III machinery in this process depended critically on the localization of Lem2-Nur1 to NE ‘tails’. We conclude that Lem2-Nur1 must be first released by the ESCRT-III/Vps4 machinery from heterochromatin during interphase to be able to enrich at the NE sealing sites in order to re-establish nucleocytoplasmic compartmentalisation.

Our findings define a mechanism through which the ESCRT-III/Vps4 machinery controls tethering of heterochromatin to the Lem2-Nur1 complexes during interphase. We propose the following model (Fig. 4j): (1) the Lem2-Nur1 complex binds to heterochromatin; (2) Cmp7 recognises this bound state and interacts with Lem2; (3) Cmp7 then recruits Vps32 and Vps4, which (4) work together to catalyse the release of Cmp7 from Lem2. We speculate that this remodelling step (5) concurrently promotes the detachment of Lem2 from heterochromatin, perhaps by modulating the interaction between Lem2 and Nur1. Repeated cycles of binding and release from heterochromatin would allow Lem2 and Nur1 to diffuse laterally along the INM. Mitotic signalling promotes bulk release of heterochromatin from Lem2-Nur1 complexes and their eventual enrichment at ‘tail’ structures. At this stage, Lem2-Nur1 complexes again engage ESCRT-III/Vps4 to promote re-establishment of nucleocytoplasmic compartmentalisation, still in the presence of the spindle. This could occur through stabilising the ‘tail’ structure and effectively sealing the membrane onto the spindle. Indeed, recent work suggests that in human cells Lem2-Cmp7 assemblies may act as a molecular sealant aiding nuclear membrane attachment to spindle microtubules^43^. Of note, we show that Lem2 requires Cmp7 to enrich at NE ‘tails’ and establish nucleocytoplasmic compartmentalization (Fig. 4h).

The release of chromosomes from the NE, whether through breakdown of the NE or dissociation from an intact membrane, is a common feature of mitosis in eukaryotes^38–41^. Indeed, in cultured human cells, the artificial tethering of histones to ER membranes leads to chromatin bridges and failure in chromosome segregation and chromatin organisation^44^. Our data indicate that ESCRT-III/AAA-ATPase Vps4 functions to release heterochromatin from the INM and this active process is critical for normal progression through mitosis. It will be interesting to see if Vps4-assisted INM-chromatin remodelling contributes to post-mitotic reformation of the NE in metazoan cells, where membrane spreads around the genome through a Brownian ratchet-like mechanism. Moreover, the dynamicity of chromatin-NE interactions afforded by ESCRT-III/Vps4 may provide a yet to be explored regulation modality shaping interphase genome organisation and function.

## Acknowledgements

We are grateful for Y. Gu, M. Makarova and R. Mori for discussions, E. Makeyev for the feedback on the manuscript, and M. Balasubramanian for an independently constructed *S. pombe vps4∆* strain. G.H.P was in part supported by a Boehringer Ingelheim Fonds PhD fellowship (2016-2018). This work was supported by the Austrian Science Fund grants to D. T. (FWF-Y444-B12, P30263, P29583) and MCBO (W1101-B18) and the Wellcome Trust Senior Investigator Award (103741/Z/14/Z) to S. O.

## Materials and Methods

### Method details

#### Culturing of yeast strains

Standard fission yeast methods and media were used^45–48^. *S. japonicus* and *S. pombe* cells were typically maintained on yeast extract with supplements (YES) rich medium 2% agar plates at 30°C. For experiments, cells were grown to the exponential growth phase (OD_595 ~_ 0.2-0.4) in liquid YES in baffled flasks in a shaker incubator at 30°C, at 200 RPM. *S. japonicus* cells were mated on SPA and dissected using a micro-dissector (Singer Instruments).

#### Generation of fission yeast mutants

All tagged proteins were tagged at their endogenous loci, with expression driven by the endogenous promoters, except when noted. Partial open reading frames (ORFs) and regions downstream of genes of interest were cloned into pJK210-based plasmids containing either mCherry (pSO730 for *S. j*) or eGFP (pSO729 for *S. j*; pSO32 for *S. p*) and the full-length *S. japonicus* or *S. pombe ura4*^+^ gene. The *S. japonicus* Lem2-GFP construct was cloned into a pJK210-based plasmid carrying the *kanMX* resistance cassette (pSO820). Gene deletions were obtained using a plasmid-based or a PCR-based method. Targeting constructs were made by cloning regions flanking the gene of interest into pJK210-based plasmids containing *kanR* (pSO820 for *S. j*) or *natR* (pSO893 for *S. j*) resistance cassettes, or the respective *ura4^+^* genes (pSO550 for *S. j*; pSO13 for *S. p*). PCR-based knockouts were obtained by amplifying the *hygR* resistance cassette flanked by 80 base pairs flanking the target gene. Plasmids were linearized before transformation. Transformation of *S. japonicus* was done by electroporation^46^. *S. pombe* cells were transformed using lithium acetate and heat shock^47^. Selection was performed on YES agar plates containing G418 (Sigma), nourseothricin (HKI Jena), hygromycin B (Roche) or minimal media (EMM) agar plates lacking uracil.

#### Generation of Vps24, Vps4 and *Cmp7* functional tagged constructs

The following constructs were designed with restriction sites flanking the 3’overlap region, endogenous promoter-ORF, tag and terminator: XhoI-3’UTR-SrfI-promoter-Vps4-AscI-3xHA-eGFP-PacI-terminator-SacII, XhoI-3’UTR-EcoRV-promoter-Vps24-AscI-LAP-eGFP-PacI-terminator-XbaI, EcoRV-3’UTR-SrfI-promoter-Cmp7-AscI-LAP-eGFP-PacI-terminator-XbaI. Constructs were synthesised by Eurofins. The eGFP of the Cmp7 construct was switched to mNeonGreen using Gibson assembly (NEB). Each construct was cloned into a pJK210-based vector for transformation into *S. japonicus*. Plasmids were linearized with SrfI or EcoRV respectively and transformed into yeast, replacing the endogenous allele. The *vps4*^*EQ*^ and *vps4*^*ts*^ alleles were constructed by site-directed mutagenesis PCR based on corresponding budding yeast mutations^22^.

#### Canavanine sensitivity assay for functionality of ESCRT-III/Vps4 constructs

ESCRT mutants show inhibited growth in the presence of the arginine analogue canavanine^49^. We titrated canavanine concentration in EMM plates with supplements from 0-10 µg ml^-1^ and determined the optimal working concentration for *S. japonicus* to be around the 4-4.5 µg ml^-1^ level (data not shown). Note that this is higher than the working concentration for *S. cerevisiae*. *vps4-3xHA-GFP* and *vps24-LAP-GFP* strains grew similar to WT when exposed to 4.5 µg ml^-1^ canavanine (Supplemental Fig. 2a). This suggests that the MVB pathway and therefore the ESCRT machinery is functional in these cells.

#### FM4-64 staining

Cells were grown to early exponential growth phase. 10 ml of cells were spun down at 3000 RPM for 1 min and resuspended in 100 µl YES media. 1 µl of FM4-64 (Thermo Fisher, stock 1 mg/ml in DMSO) was added and incubated with cells for 5 min at 30°C. Subsequently, the cells were washed twice with YES and resuspended in 10 ml of YES and grown for 1 hr at 30°C in a shaking incubator before imaging.

#### Vps4 inactivation using a temperature-sensitive allele

Cells were grown in liquid culture overnight at 25°C in a shaking incubator to early exponential growth phase. The culture was shifted to 36°C and samples were taken at time 0 and then at 1 and 4 hours. Slides for imaging were prepared on a heat block at 36°C. Imaging was performed in a pre-heated chamber at 36°C.

#### Colony-forming unit (CFU) assay

Cells were grown overnight at 30°C in a shaking incubator. The next day, cells with an OD_595_ of 0.2-0.3 were diluted to 0.1. Next, three serial dilutions of 10x, 100x and 1000x were prepared. Of these dilutions, 50 µl of cells were plated in triplicate on YES plates and spread using glass beads. After two days of incubation at 30°C the plates were scanned using an Epson Perfection V700 Photo scanner and Epson Scan software. CFU count and size were automatically analysed using OpenCFU software^50^.

#### Microscopy

For imaging, cells grown in liquid YES media to early exponential growth phase were placed on a thin YES 2% agarose strip and immobilised by a cover slip, which was sealed with wax. This slide was first rested for 30 min at the 30°C. During imaging cells where kept at 30°C in an environmental control chamber. Imaging was performed on a Nikon Eclipse Ti-E inverted system equipped with CSU-X1 spinning disk confocal unit and 600 series SS 488nm, SS 561nm lasers. Images were obtained with an Andor iXon Ultra U3-888-BV monochrome EMCCD camera using a Nikon CFI Plan Apo Lambda 100x/1.45NA objective lens. Images presented in this report are Z-projections unless noted otherwise. For microscopy images contrast and brightness were adjusted for each individual image for optimal visibility unless noted otherwise. For time-lapse imaging, laser power and exposure time were adjusted to minimise photobleaching.

#### Protein expression and purification

Expression of proteins was performed as previously described^21,51^. Recombinant proteins were expressed in C41(DE3) pLysS *E. coli* (Lucigen) and induced at 37°C for 4h in 1mM IPTG (Thermo R0392). GST-tagged proteins were purified with Glutathione Sepharose 4B (GE Healthcare, GE17-0756-01), washed and either eluted with glutathione of cleaved with PreScission protease (GE Healthcare, 27084301) overnight at 4°C. Recombinant proteins were dialyzed in a Slide-a-lyzer (VWR, 514-0172) overnight in ATPase buffer (100mM potassium acetate, 5mM magnesium acetate, 20mM Hepes 7.4) and were additionally purified via a Superdex 2000 column (GE Healthcare). Proteins were flash frozen in liquid nitrogen and stored at −80°C until further usage.

#### Plasmids used for in vitro experiments

| Name | Insert | Backbone |
| --- | --- | --- |
| pSS76 | GST-Lem2<sup>564-673</sup> | pGex 6P1 |
| pSS98 | GST-Did4 | pGex 6P1 |
| pSS113 | GST-Cmp7<sup>242-436</sup> | pGex 6P1 |
| pSS114 | GST-Cmp7<sup>242-436</sup>-3xFLAG | pGex-6P1 |
| pSS115 | Vps4-3xHA | pGex 6P1 |
| pSS116 | Vps4<sup>E233Q</sup>-3xHA | pGex 6P1 |
| pSS117 | Vps32-3xMyc | pGex 6P1 |

#### GST pulldown assays

GST pulldown assays were performed as previously described ^31,52^. Pierce magnetic glutathione beads (Thermo, 78602) were incubated in 0.1%BSA in ATPase buffer overnight. 5µg of GST tagged proteins were bound to beads for 2 hours at 4°C, washed and optionally incubated with 500ng of Cmp7-3xFLAG and/or Vps32-3xMyc for 1 hour at 4°C. 500ng of Vps4^E233Q^ −3xHA or Vps4-3xHA was added in the presence or absence of 1mM ATP for 10 minutes at room temperature. After five washing steps in ATPase buffer with 600mM NaCl, proteins were eluted from beads using sample buffer (2%SDS, 100mM Tris 6.8, 10% glycerol, 5% beta-mercaptoethanol, 0.01% bromophenol blue) at 96°C for 10 minutes. Samples were separated on a 12.5% SDS Page. Proteins were either stained by Brilliant Blue Coomassie or subjected to Western blotting.

#### Antibodies and Reagents

Anti-FLAG (3165) and anti-Myc (4439) antibodies were purchased from Sigma. Anti-HA antibody was purchased from Cell Signalling (C29F4). ATP was purchased from Sigma (A-2383).

### Quantification and data analysis

#### Analysis software

All imaging data were analysed and imaging data figures were prepared in ImageJ^53,54^. Data was analysed using Microsoft Excel. Statistics was performed in Graphpad Prism. Graphs were prepared in Graphpad Prism. Figures were prepared in Adobe Indesign or Adobe Illustrator.

#### NE resealing assay

For imaging of NE resealing cells undergoing mitosis were typically imaged every 30 seconds for a maximum duration of 40 min (n=15 cells). Z-stacks of slices with 0.5 µm distance (total stack 6.0 µm) were obtained at each time point. Maximum Z-projections were made and the average intensity of a circle with a diameter of 0.695 μm (10 pixels) was measured in the nucleus next to the SPB (used as a spatial cue).

#### Analysis of recruitment of Vps4 to distal NE tails

For time-lapse imaging, cells were imaged every 1 minute for a maximum duration of 30 minutes. Z-stacks of slices with 0.5 µm distance (total stack 4.5 µm) were obtained at each time point. Maximum Z-projections were made and a line was drawn of 8 pixels wide and 80 pixels long starting from the SPB to the end of the tail structure. Fluorescence intensity was extracted using the plot-profile function for 10 cells (Fig. 2c). For Fig. 2d (timing of Vps4 recruitment to the tails) 13 cells were followed from NE rupture to spindle breakdown.

#### Recruitment of ESCRT-III/Vps4 to the NE during interphase

For imaging of persistent recruitment of Vps4-GFP to the NE (Fig. 3e and Supplemental Fig. 4c), cells were imaged every 10 seconds for 5 minutes. Z-stacks of slices with 0.5 µm distance (total stack 4.5 µm) were obtained at each time point. One replicate experiment constituted imaging several fields of view for a total period of 1 hr and was repeated three times. For analysis, single confocal slices of each cell at each time-point were screened for fluorescence signal on the NE. The ImageJ line-plot tool was used to confirm if the fluorescent signal was an ESCRT-III/Vps4 event directly on the NE marked by Nup189-mCherry. Persistent events were defined as events that lasted from the start to the end of each 5-minute time-lapse.

#### Measurement of distance between Taz1 or Mis6 and the NE

Z-stacks of slices with 0.25 µm distance (total stack 4.5 µm) were obtained. To measure the distance of a Taz1 or Mis6 focus to the NE a line was drawn in a single Z-slice from the centroid of Taz1/Mis6 signal to the centroid of the NE signal and the length in pixels was measured and amplified by 0.0695 to obtain the physical distance. In many cases the focus was nearer to the NE in Z than in Y or X. In this case the orthogonal view function of ImageJ was used to count the number of Z-steps from the centre of the focus to the centre of the NE signal as the distance (resulting in a lot of data points having 0.25, 0.5, 0.75 etc. as distance).

## Author contributions

G.H.P. designed, performed and analysed all genetics and cell biological experiments and wrote the first version of the manuscript. S.S. performed all in vitro biochemical experiments and validated the functionality of ESCRT-III/Vps4 tagged constructs. S.S. and G.H.P. constructed ESCR-III/Vps4 fluorescent protein constructs. G.H.P., S.S., D.T., S.O., conceived experiments and reviewed and edited the manuscript. D. T and S.O. supervised the project.

