## Supplementary information for "ESCRT-III/Vps4 controls heterochromatin-nuclear envelope attachments"

Contains:

Extended Data figures 1-5 page 2-10

Strain list (table 1) page 11-12

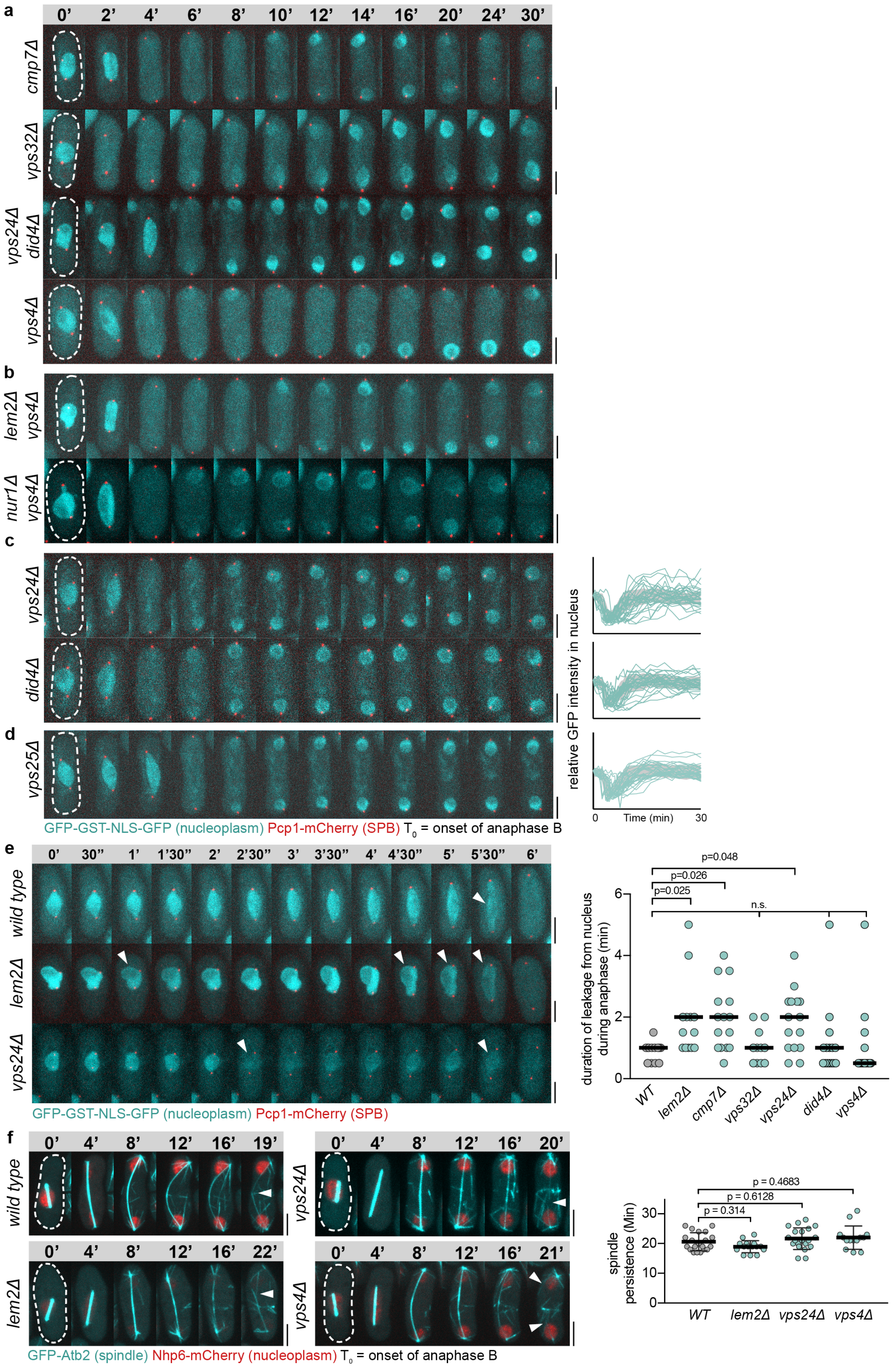


**Supplemental Fig. 1, related to Fig. 1.** (**a**) Representative GFP-NLS Pcp1-mCherry-expressing cells of indicated genotypes used for the quantification in Fig. 1g. (**b**)

Representative GFP-NLS Pcp1-mCherry-expressing cells of indicated genotypes used for the quantification in Fig. 1h. (**c**) *Left*, Representative GFP-NLS Pcp1-mCherry-expressing cells of indicated genotypes used for the quantification in resealing assays shown on the *right*. (**d**) Representative GFP-NLS Pcp1-mCherry-expressing *vps25Δ* (ESCRT-II) mutant cells used for the quantification in resealing assay shown on the *right*. (**e**) Premature loss of nuclear integrity during mitosis in *lem2* and ESCRT-III/Vps4 mutants. Arrows indicate loss of nuclear integrity. Note the single abrupt rupture in WT versus the repeated rupture and slow leaking of the nucleoplasm in the mutants. The same set-up as in Fig. 1 was used to quantify the duration of GFP-NLS leakage from the nucleus after anaphase onset. Medians are indicated. Knuskal-Wallis multiple comparison test. (**f**) Time-lapse maximum projection sequences of GFP-Atb2 and Nhp6-mCherry expressing cells of indicated genotypes starting from the onset of anaphase B. Arrows indicate the breakage of the spindle. Quantification of time between anaphase B onset and breakage of the spindle is presented on the *right* (n≥14 cells; one way ANOVA followed by Dunnett's comparison test. (**a**-**f**) Scale bars represent 5 μm.

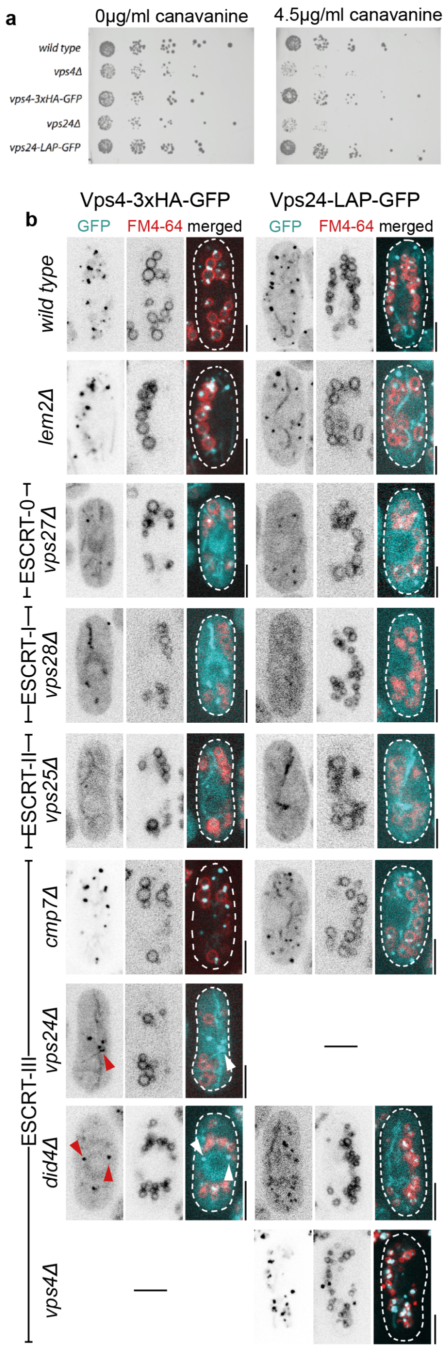


**Supplemental Fig. 2, related to Fig. 2**. (**a**) Canavanine sensitivity assay. WT, *vps4Δ*, or *vps24Δ* mutant cells or cells expressing Vps4-GFP or Vps24-GFP were spotted on EMM plates containing either 0 or 4.5 μgr ml$^{-1}$ canavanine. Note that ESCRT-III tagging does not induce canavanine sensitivity. (**b**) Shown are single confocal slices of cells expressing Vps4-GFP or Vps24-GFP that were incubated with FM4-64 for 5 minutes followed by washout of the drug and recovery for 1 hour before imaging. Note that in early ESCRT mutants Vps4 and Vps24 are no longer recruited to FM4-64 marked structures. Arrows indicate bright Vps4 foci on the NE in *vps24Δ* and *did4Δ* mutants. Scale bars represent 5 μm.

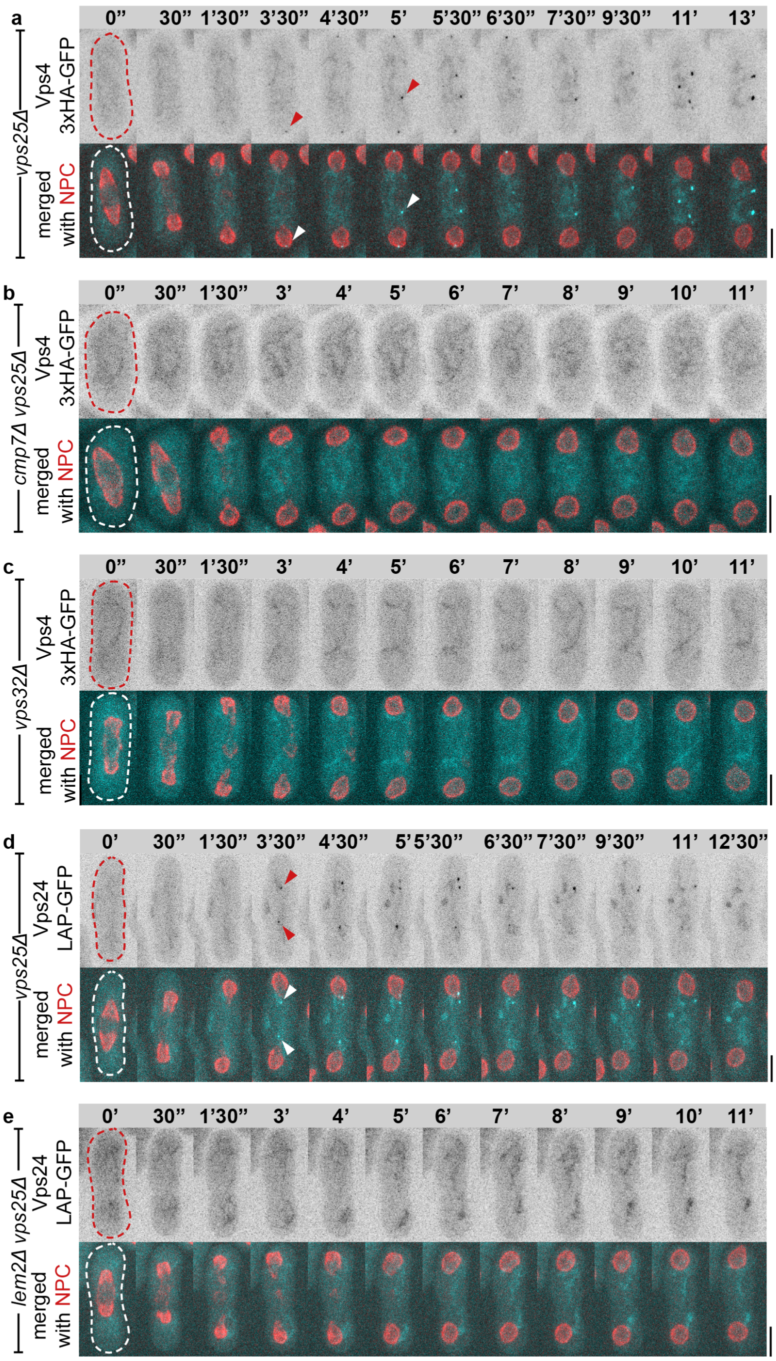


**Supplemental Fig. 3, relates to Fig. 2.** (**a**) A time-lapse sequence of a *vps25Δ* Vps4-GFP Nup189-mCherry cell starting prior to NE rupture (n=12). Note that the NPCs are excluded from the ‘tails’. (**b**) A time-lapse sequence of a *cmp7Δ vps25Δ* Vps4-GFP Nup189-mCherry cell starting prior to NE rupture (n=15). (**c**) A time-lapse sequence of a *vps32Δ* Vps4-GFP Nup189-mCherry cell starting prior to NE rupture (n=12). (**d**) A time-lapse sequence of a *vps25Δ* Vps24-GFP Nup189-mCherry cell starting prior to NE rupture (n=12). Note that Vps24 also localizes to the ‘tails’ despite not being required for re-establishment of nucleocytoplasmic compartmentalisation. (**e**) A time-lapse sequence of a *lem2Δ vps25Δ* Vps24-GFP Nup189-mCherry cell starting prior to NE rupture (n=8) shows the dependency of Vps24 on Lem2 for NE recruitment. (**a**-**e**) Shown are Z-projections of spinning disk confocal stacks, unless indicated otherwise. Scale bars represent 5 μm.

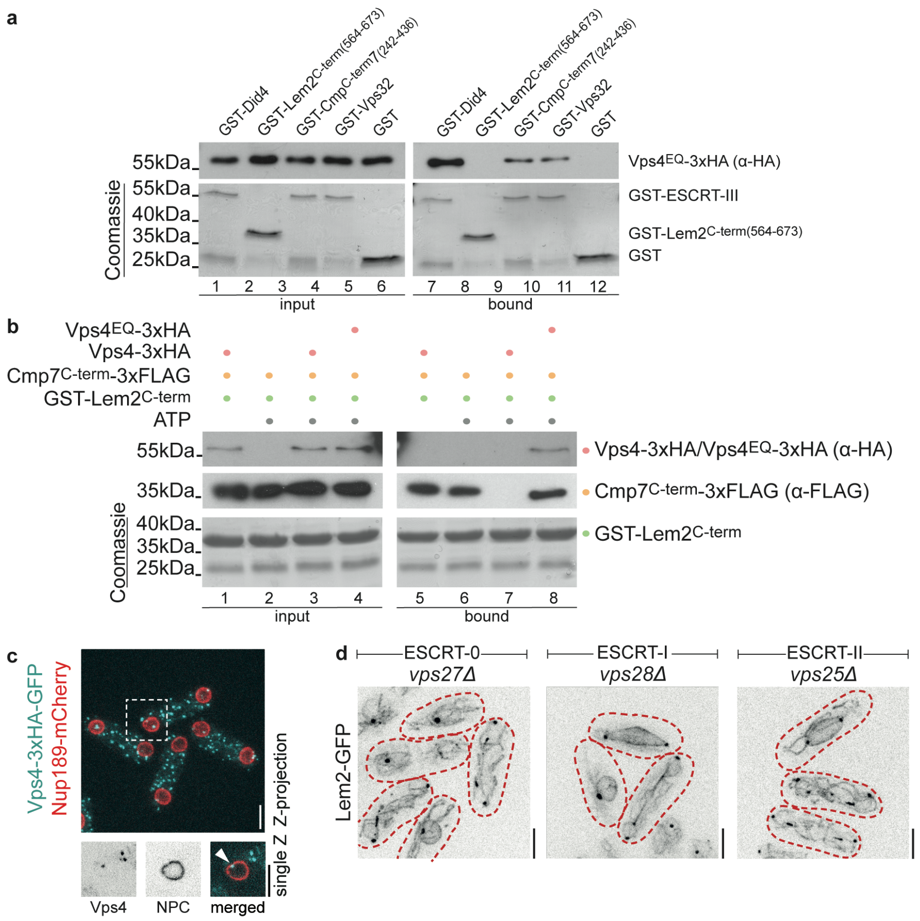


**Supplemental Fig. 4, related to Fig. 3.** (**a**) *In vitro* binding experiment. An ATPase dead Vps4$^{EQ}$ that can bind to its substrates but cannot hydrolyse ATP was added to glutathione beads coated with GST-Did4, GST-Lem2$^{Cterm}$, GST-Cmp7$^{Cterm}$ or GST-Vps32. Vps4$^{EQ}$ bound strongly to Did4 as expected and weakly to Cmp7$^{C\text{-}term}$ and Vps32. No binding of Vps4$^{EQ}$ to Lem2$^{C\text{-}term}$ could be detected. (**b**) *An in vitro* binding experiment using a setup similar to Fig. 3a. Lem2-Cmp7 complexes were incubated with either WT Vps4 or Vps4$^{EQ}$, in the presence or absence of ATP. WT Vps4 could only disassemble the Lem2-Cmp7 complex in the presence of ATP (lane 7), but not when ATP was absent (lane 5). (**c**) A Z-projected spinning disk confocal image of cells expressing Vps4-GFP and Nup189-mCherry. A magnified image below focusses on a nucleus exhibiting a persistent Vps4 focus on the NE. (**d**) Maximum projection images of spinning disk confocal stacks of cells of indicated genotypes expressing Lem2-GFP. Note that Lem2 does not cluster at the NE in 'early' ESCRT mutants. (**c**, **d**) Scale bars represent 5 μm.

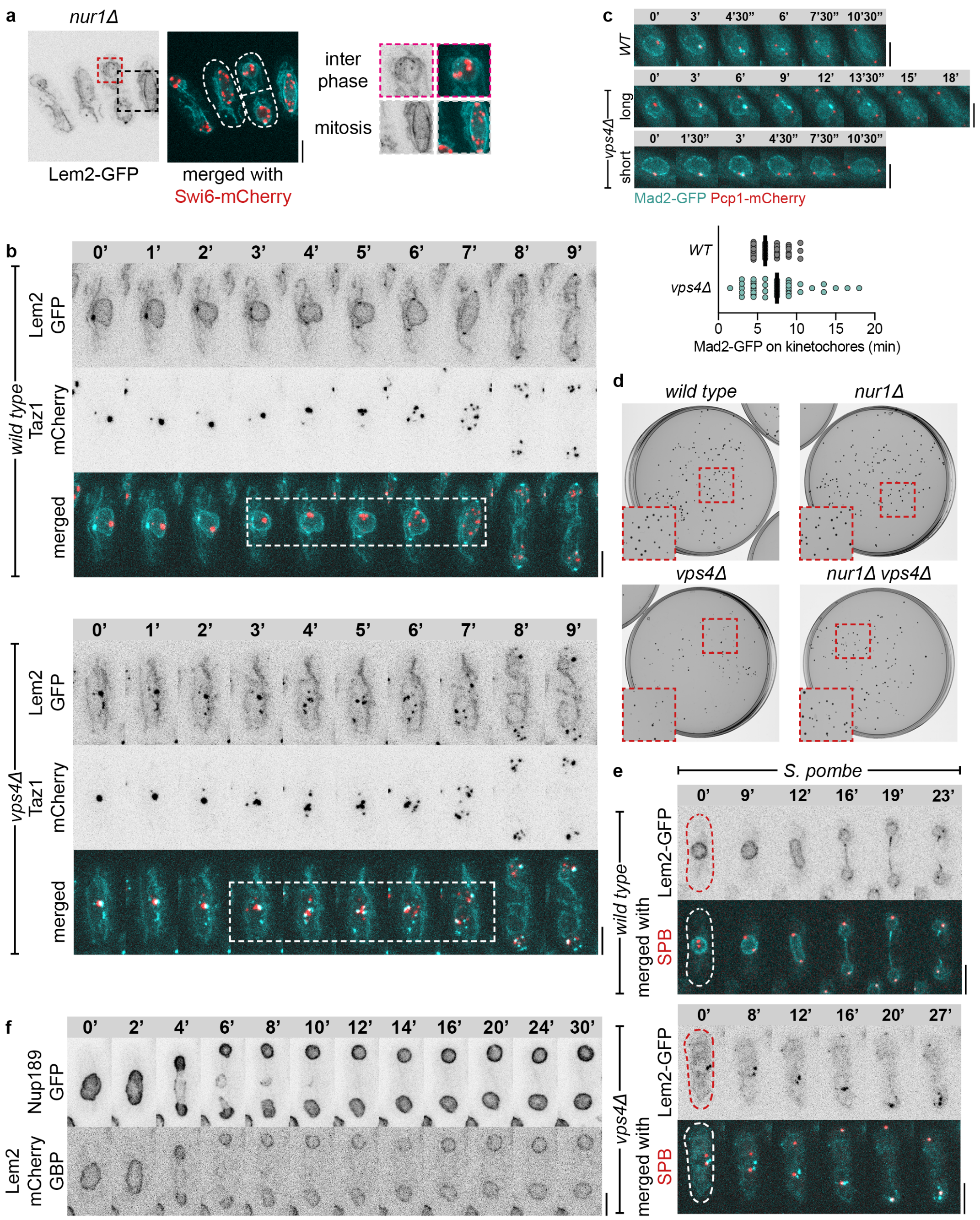


**Supplemental Fig. 5, related to Fig. 4.** (**a**) A maximum projection of a spinning disk confocal stack of *nur1Δ* cells expressing Lem2-GFP and Swi6-mCherry, with the layout as in Fig. 4a. (**b**) Full time courses and images of entire cells presented in Fig. 4c. (**c**) Time-lapse maximum projection sequences of representative cells of indicated genotypes expressing Mad2-GFP and Pcp1-mCherry (*top*). A graph quantifying the duration of Mad2-GFP presence at kinetochores in WT (n=40 cells) and *vps4Δ* (n=41 cells) (*bottom*). (**d**) Scans of plates of a single technical replicate in a CFU assay presented in Fig. 4 f-g. *Insets*, magnified images of the indicated areas. (**e**) Maximum projections of time-lapse spinning disk confocal stacks of *S. pombe* cells of indicated genotypes expressing Lem2-GFP and Pcp1-mCherry. Note that Lem2 clusters do not disassemble in mitotic *vps4Δ* cells (n≥7 cells). (**f**) Time-lapse sequence of a cell expressing Lem2-mCherry-GBP and Nup189-GFP. Note that the Lem2 signal completely overlaps with that of Nup189, indicating that it is stably associating with NPCs throughout the cell-cycle.

### Table 1: strain list

| | | |
|---|---|---|
| ***S. japonicus* strains** | | |
| All strains derived from: | | |
| SOJ11 | *matsj-P2028 ura4sj-D3 ade6sj-domE* | h- |
| SOJ88 | *matsj-P2028 ura4sj-D3 ade6sj-domE* | h+ |
| ***S. japonicus* strains constructed in this work** | | |
| SOJ88 | *matsj-P2028 ura4sj-D3 ade6sj-domE* | |
| SOJ359 | *ura4::GFP-atb2 nhp6-mCherry:ura4* | h+ |
| SOJ2001 | *ura4::GFP-GST-NLS-GFP pcp1-mCherry:ura4:kanR* | h+ |
| SOJ2003 | *lem2Δ:kanR ura4::GFP-GST-NLS-GFP pcp1-mCherry:ura4:kanR* | |
| SOJ2251 | *did4Δ:kanR* | h- |
| SOJ2370 | *cmp7Δ:kanR* | h- |
| SOJ2664 | *did4Δ:kanR ura4::GFP-GST-NLS-GFP pcp1-mCherry:ura4:kanR* | h+ |
| SOJ2668 | *cmp7Δ:kanR ura4::GFP-GST-NLS-GFP pcp1-mCherry:ura4:kanR* | h+ |
| SOJ2748 | *cmp7Δ:kanR lem2-GFP:kanR pcp1-mCherry:ura4:kanR* | |
| SOJ2775 | *vps24Δ:kanR* | h+ |
| SOJ2778 | *vps20Δ:kanR* | h+ |
| SOJ2782 | *vps4Δ:kanR* | h+ |
| SOJ2788 | *vps4Δ:kanR ura4::GFP-GST-NLS-GFP pcp1-mCherry:ura4:kanR* | h- |
| SOJ2791 | *vps24Δ:kanR ura4::GFP-GST-NLS-GFP pcp1-mCherry:ura4:kanR* | h- |
| SOJ2841 | *vps24Δ:kanR lem2-GFP:kanR pcp1-mCherry:ura4:kanR* | |
| SOJ2842 | *vps4Δ:kanR lem2-GFP:kanR pcp1-mCherry:ura4:kanR* | |
| SOJ2843 | *lem2Δ:natR* | h+ |
| SOJ2845 | *lem2-GFP:kanR taz1-mCherry:ura4* | h- |
| SOJ2863 | *lem2-GFP:kanR pcp1-mCherry:ura4:kanR* | |
| SOJ2867 | *vps4Δ:kanR lem2-GFP:kanR taz1-mCherry:ura4* | |
| SOJ2883 | *vps25Δ:kanR* | h+ |
| SOJ2885 | *vps27Δ:kanR* | h+ |
| SOJ2890 | *vps28Δ:kanR* | h+ |
| SOJ2895 | *vps27Δ:kanR lem2-GFP:kanR* | |
| SOJ2896 | *vps25Δ:kanR lem2-GFP:kanR* | |
| SOJ2927 | *vps28Δ:kanR lem2-GFP:kanR* | |
| SOJ2976 | *lem2-GFP:kanR swi6-mCherry:ura4* | |
| SOJ2977 | *vps4Δ:kanR lem2-GFP:kanR swi6-mCherry:ura4* | |
| SOJ2981 | *vps4-3xHA-GFP:ura4* | h+ |
| SOJ2982 | *lem2Δ:natR vps4-3xHA-GFP:ura4* | h+ |
| SOJ2983 | *cmp7Δ:kanR vps4-3xHA-GFP:ura4* | h+ |
| SOJ2984 | *vps24Δ:kanR vps4-3xHA-GFP:ura4* | h+ |
| SOJ3000 | *nur1-mCherry:ura4 lem2Δ:natR* | |
| SOJ3018 | *vps24-LAP-GFP:kanR* | h+ |
| SOJ3034 | *vps25Δ:kanR vps4-3xHA-GFP:ura4* | h+ |
| SOJ3057 | *vps4-3xHA-GFP:ura4 nup189-mCherry:ura4* | h- |
| SOJ3062 | *vps25Δ:kanR vps24-LAP-GFP:kanR nup189-mCherry:ura4* | h+ |
| SOJ3065 | *lem2-GFP:kanR nur1-mCherry:ura4* | h- |
| SOJ3079 | *vps27Δ:kanR vps24-LAP-GFP:ura4* | |
| SOJ3080 | *vps27Δ:kanR vps4-3xHA-GFP:ura4* | |
| SOJ3081 | *vps28Δ:kanR vps4-3xHA-GFP:ura4* | |
| SOJ3097 | *did4Δ:kanR vps4-3xHA-GFP:ura4* | |
| SOJ3099 | *did4Δ:kanR vps24-LAP-GFP:ura4* | |
| SOJ3132 | *cmp7Δ:kanR vps24-LAP-GFP:ura4* | h+ |
| SOJ3233 | *vps25Δ:kanR vps4-3xHA-GFP:ura4 nup189-mCherry:ura4* | h- |
| SOJ3237 | *lem2Δ:natR vps25Δ:kanR vps4-3xHA-GFP:ura4 lem2-mCherry:ura4* | h+ |
| SOJ3244 | *lem2Δ:natR vps25Δ:kanR vps24-LAP-GFP:ura4 nup189-mCherry:ura4* | |
| SOJ3263 | *vps4-3xHA-GFP:ura4 lem2-mCherry:ura4* | |
| SOJ3267 | *lem2Δ:natR vps24-LAP-GFP:ura4* | |

| | | |
|---|---|---|
| SOJ3268 | *vps25Δ:kanR vps24-LAP-GFP:ura4* | |
| SOJ3284 | *lem2-GFP:kanR ura4::adh1pro-GST-NLS-mCherry* | |
| SOJ3291 | *vps4(E233Q)-3xHA-GFP:kanR lem2-mCherry* | h+ |
| SOJ3293 | *vps28Δ:kanR vps24-LAP-GFP:ura4* | |
| SOJ3294 | *vps4Δ:kanR vps24-LAP-GFP:ura4* | |
| SOJ3305 | *did4Δ:kanR vps4-3xHA-GFP:ura4 lem2-mCherry* | |
| SOJ3322 | *vps25Δ:kanR ura4::GFP-GST-NLS-GFP pcp1-mCherry:ura4:kanR* | |
| SOJ3341 | *vps24Δ:kanR vps4-3xHA-GFP:ura4 lem2-mCherry* | |
| SOJ3355 | *lem2Δ:natR vps4Δ:natR ura4::GFP-GST-NLS-GFP pcp1-mCherry:ura4:kanR* | |
| SOJ3444 | *vps24Δ:kanR lem2-GFP:kanR swi6-mCherry:ura4* | h- |
| SOJ3477 | *vps4Δ:natR ura4::GFP-atb2 nhp6-mCherry:ura4* | |
| SOJ3478 | *vps24Δ:natR ura4::GFP-atb2 nhp6-mCherry:ura4* | |
| SOJ3481 | *lem2Δ:natR ura4::GFP-atb2 nhp6-mCherry:ura4* | |
| SOJ3500 | *cmp7Δ:kanR lem2-GFP:kanR swi6-mCherry:ura4* | |
| SOJ3514 | *cmp7Δ:kanR vps25Δ:kanR vps4-3xHA-GFP:ura4 lem2-mCherry:ura4* | h- |
| SOJ3578 | *taz1-GFP:ura4 nup189-mCherry:ura4 pcp1-mCherry:ura4:kanR* | |
| SOJ3592 | *vps24Δ:hygR did4Δ:kanR ura4::GFP-GST-NLS-GFP pcp1-mCherry:ura4:kanR* | h+ |
| SOJ3596 | *nur1Δ:hygR* | h- |
| SOJ3623 | *mad2-GFP:ura4 pcp1-mChery:ura4:kanR* | h- |
| SOJ3638 | *mis6-GFP:ura4 nup189-mCherry:ura4 pcp1-mCherry:ura4:kanR* | h+ |
| SOJ3640 | *vps4Δ:kanR taz1-GFP:ura4 nup189-mCherry:ura4 pcp1-mCherry:ura4:kanR* | |
| SOJ3650 | *vps4Δ:kanR mad2-GFP:ura4 pcp1-mChery:ura4:kanR* | |
| SOJ3697 | *vps32Δ:hygR ura4::GFP-GST-NLS-GFP pcp1-mCherry:ura4:kanR* | |
| SOJ3700 | *vps32Δ:kanR lem2-GFP:kanR pcp1-mCherry:ura4:kanR* | |
| SOJ3701 | *vps32Δ:kanR lem2-GFP:kanR swi6-mCherry:ura4* | |
| SOJ3727 | *vps4ts(I307T,L327S)-3xHA-GFP:ura4 lem2-mCherry:ura4* | h+ |
| SOJ3728 | *vps4Δ:kanR mis6-GFP:ura4 nup189-mCherry:ura4 pcp1-mCherry:ura4:kanR* | |
| SOJ3748 | *cmp7-LAP-mNeonGreen:kanR lem2-mCherry:ura4* | h+ |
| SOJ3756 | *nur1Δ:hygR vps4Δ:kanR lem2-GFP:kanR swi6-mCherry:ura4* | |
| SOJ3762 | *vps32Δ:hygR vps4-3xHA-GFP:ura4 nup189-mCherry:ura4* | |
| SOJ3764 | *nur1Δ:hygR lem2-GFP:kanR swi6-mCherry:ura4* | |
| SOJ3765 | *nur1Δ:hygR vps4Δ:kanR* | |
| SOJ3770 | *nur1Δ:hygR ura4::GFP-GST-NLS-GFP pcp1-mCherry:ura4:kanR* | h+ |
| SOJ3799 | *vps4Δ:kanR cmp7-LAP-mNeonGreen:kanR lem2-mCherry:ura4* | |
| SOJ3800 | *vps4Δ:kanR nur1-mCherry:ura4 lem2-GFP:kanR* | |
| SOJ3803 | *nur1Δ:hygR vps4Δ:kanR ura4::GFP-GST-NLS-GFP pcp1-mCherry:ura4:kanR* | |
| SOJ3812 | *nur1Δ:hygR vps25Δ:kanR vps4-3xHA-GFP:ura4 nup189-mCherry* | |
| SOJ3940 | *vps25Δ:kanR vps4-3xHA-GFP:ura4 lem2-mCherry:ura4* | h- |
| SOJ3989 | *lem2-mCherry-TEV-GBP:ura4 nup189-GFP:ura4 ura4::atb2pro:NLS-GST-mCherry* | |
| SOJ3990 | *lem2-mCherry-TEV-GBP:ura4 nup189-GFP:ura4 ura4::atb2pro:NLS-GST-mCherry* | |
| SOJ3992 | *nup189-GFP:ura4 lem2-mCherry-TEV-GBP:ura4* | |
| SOJ4001 | *lem2-mCherry-TEV-GBP:ura4 nup189-GFP:ura4 ura4::atb2pro:NLS-GST-mCherry* | |

| ***S. pombe* strains** | | |
|---|---|---|
| All strains derived from: | | |
| SO1082 | *ade6-M210 ura4-D18 leu1-32* | h- |
| SO1083 | *ade6-M210 ura4-D18 leu1-32* | h+ |
| SO2865 | *ade6-210 ura4-D18 leu1-32* | h+ |
| SO2866 | *ade6-216 ura4-D18 leu1-32* | h- |
| ***S. pombe* strains constructed in this work** | | |
| SO8284 | *lem2-GFP:ura4 pcp1-mCherry:ura4* | h- |
| SO8355 | *vps4Δ:ura4 lem2-GFP:ura4 pcp1-mCherry:ura4* | |
